# Heightened Sensitivity to Voice Loudness Changes in Parkinson’s Disease

**DOI:** 10.1101/2025.05.29.656769

**Authors:** Francisco Contreras-Ruston, Antonio Criscuolo, Antonio Callén, David Cucurell, Jordi Navarra, Sonja A. Kotz

## Abstract

Parkinson’s disease (PD) affects voice and speech production, often resulting in reduced vocal intensity and monotonous speech. Recent studies have suggested that these changes can be partially explained by altered sensory feedback processing when producing speech. Individuals with PD (IwPD) may fail to monitor sensory feedback from their own voice, impairing their ability to adjust voice and speech when sensory input differs from expectations. In this study, we investigated sensory feedback processing in PD by looking at the sensory attenuation typically observed in event-related responses (ERP) to the self-generated voice. When sensory feedback processing is intact, the P50, N100, and P200 ERP responses to the self-generated voice are more attenuated than to an externally-generated voice. Twenty-three IwPD and 23 healthy controls (HCs) participated in a voice playback study that comprised three conditions: self-generated voice (auditory-motor condition; AMC), externally-generated voice (auditory-only condition; AOC), and motor-only. The AMC and AOC conditions also included an amplitude modulation of the voice (0/+15dB). Linear mixed models assessed group differences in ERP morphology. While groups did not differ in their P50 and P200 responses, there was a significant group–condition– loudness interaction for the N100. Follow-up analyses showed that IwPD displayed much larger N100 error responses for unexpected loudness modulations as compared to HC. This observation suggests that IwPD may process voice modulations differently than HC. The hypersensitivity to loudness changes may underlie IwPD’s difficulties in processing and adapting their voice acoustics.

## 1. INTRODUCTION

Parkinson’s disease (PD) is a progressive neurodegenerative disorder characterized by motor symptoms such as tremors, rigidity, bradykinesia, and postural difficulties (Poewe *et al*., 2017). Next to motor symptoms, individuals with Parkinson’s disease (IwPD) also display non-motor symptoms, including sensory alterations, that can affect the integration of auditory and proprioceptive feedback. Such compromised sensory feedback processing can affect voice and speech production, manifesting as reduced vocal loudness and imprecise articulation (Ma *et al*., 2020). In particular, vocal production in IwPD is often marked by hypokinetic dysarthria, which affects voice modulation and articulation and negatively impacts communication and quality of life (Ma *et al*., 2020).

Voice and speech production involves a sophisticated, multi-layered process that relies on the precise interaction between motor planning, motor execution, sensory feedback (i.e., auditory and somatosensory), and adaptation to the acoustic characteristics in the environment (e.g., noise; Blakemore et al., 2000; Knolle et al., 2019; Whitford, 2019). Auditory and somatosensory integration engage cortico-subcortical circuits (including the auditory and motor cortices and the basal ganglia), which play a key role in sensory feedback processing (Nagy et al., 2006; Schwartze & Kotz, 2016, 2024; Cardenas et al., 2020). Recent evidence also highlighted the contribution of subcortical structures, such as the brainstem, in early auditory processing. For instance, Mollaei et al. (2022) used frequency-following responses (FFR) to show that IwPD display enhanced encoding of the fundamental frequency (F0) in the brainstem. This finding suggests that dopaminergic dysfunction in PD may affect early auditory representations, potentially altering bottom-up sensory signals that guide vocal motor control (Mollaei *et al*., 2022). Voice production relies on two complementary processes. First, an adaptive control system integrates motor commands with sensory feedback and environmental cues to regulate vocal loudness; for example, speakers unconsciously increase their loudness in noisy environments by monitoring and adjusting their sensory feedback (Hilger *et al*., 2022). Second, sensory attenuation allows distinguishing between self-generated and externally-generated sounds, and predictive (forward) models generate an internal estimate of the expected voice signal, which is then compared to the actual auditory input. This comparison leads to the suppression of one’s own voice and prevents attentional overload of the auditory system (Knolle *et al*., 2019). These two processes, real-time voice adjustment and predictive sensory suppression, support precise voice output and consistent auditory feedback. However, changes in sensory feedback processing can compromise the regulation of vocal loudness and articulatory precision in PD (Huang *et al*., 2016).

If impaired sensory feedback processing disrupts the modulation of vocal loudness and articulatory precision in IwPD, this might show common voice symptoms, such as hypophonia (Liu *et al*., 2012; Railo *et al*., 2020). This symptom results from disruptions in neural circuitry involved in distinguishing between expected and actual sensory feedback. For instance, altered functional connectivity between the basal ganglia and periaqueductal gray regions, which is critical for motor-sensory integration, may underlie the reduced voice loudness observed in IwPD (Rektorova *et al*., 2012). Importantly, feedforward projections responsible for expected acoustic features such as loudness and adaptation of actual sensory feedback when expectations are not met, seem both affected in IwPD (Abur *et al*., 2021). Several studies have indicated that IwPD have difficulties in appropriately anticipating and adapting the acoustic features of their own voices (Liu *et al*., 2012; Abur *et al*., 2018). Notably, it was shown that while IwPD can display exaggerated compensatory responses to pitch perturbations, their responses to formant shifts can be markedly reduced. This dissociation suggests that IwPD may differentially weigh acoustic cues in voice motor control, possibly relying more heavily on pitch-related feedback while underutilizing articulatory-specific spectral information (Mollaei et al., 2016), and that similar imbalances may extend to loudness cues, an acoustic feature that remains understudied.

Clinically, this would manifest as difficulties in accurately perceiving one’s own vocal loudness, leading to an overestimation of loudness levels due to impaired sensory feedback processing (Clark *et al*., 2014). Given that these rapid predictive and corrective voice adaptations typically occur within less than 300 ms (Behroozmand et al., 2016), event-related potentials (ERPs) offer an effective method to investigate these early stages of sensory feedback processing in the PD voice.

ERPs are sensitive in capturing rapid sensory feedback processing, as they show in millisecond resolution. ERPs such as the P50, N100, and P200 components are indicators of sensory and sensory feedback processing in voice production. For example, the P50 is linked to sensory gating, which is essential for filtering out redundant or irrelevant auditory stimuli (Brinkmann *et al*., 2021). An increased P50 amplitude in IwPD was interpreted as reduced sensory habituation that is expected in sensory gating (De Groote *et al*., 2020; Hassin-Baer *et al*., 2022).

The N100 has been related to the match/mismatch between expected and actual sensory feedback. The match/mismatch effect shows in a reduced/enhanced N100 response in self-voice production (Heinks-Maldonado *et al*., 2005; Knolle *et al*., 2019). However, there is currently very little evidence on the N100 suppression effect in IwPD. Studies that explored the ascribed match/mismatch effect for other effectors in IwPD, such as postural adjustments or adaptation in motor tasks (Olson *et al*., 2019; Palmisano *et al*., 2020), argued that similar deficits could be observed in the PD voice and speech production (Huang *et al*., 2016). These results would explain why brain circuitry that normally optimizes sensory feedback processing and suppresses neural responses to expected sensory input may be affected in IwPD across multiple motor effectors (see Puyjarinet et al., 2019). In healthy individuals, the N100 suppression effect reflects the precision of monitoring expected and actual sensory feedback. In the case of a mismatch, an error that minimizes the suppression effect is elicited (see Knolle et al., 2019). As dopamine might regulate error processing and is depleted in PD, this might lead to less accurate monitoring of errors due to impaired sensory feedback processing in voice production (Huang et al., 2016, 2019). In fact some prior ERP evidence in IwPD reported a reduced N100 suppression effect during vocalization, suggesting a deficit in speech monitoring (Railo et al., 2020). However, such evidence is sparse and has predominantly been explored in pitch modulation, which does not differ between IwPD and HCs (Huang *et al*., 2016). Thus, the question of sensory feedback processing in the context of voice (loudness) adaptation remains an open question in IwPD.

The last component of interest, the P200 has been related to auditory attention and sensory feedback processing, and the regulation of voice control (Behroozmand et al., 2009; Duggirala et al., 2023). In IwPD, an increased P200 amplitude has been interpreted as enhanced neural responsiveness and a potential compensatory strategy in response to errors in sensory feedback processing (Huang et al., 2016; Railo et al., 2020). Hence, in IwPD, an enhanced P200 response might indicate greater dependence on sensory feedback to regulate voice production (Garrido-Vásquez *et al*., 2013; Li *et al*., 2021). This modulation was also associated with attention to sensory feedback, suggesting that changes in P200 amplitude could reflect a reduction in attentional resources devoted to error processing, implying a reduced ability to detect auditory discrepancies (Behroozmand *et al*., 2009), or conversely, an increased allocation of attention as a compensatory mechanism in response to sensory error detection (Duggirala *et al*., 2023).

Despite this evidence, existing ERP data in PD voice production remains limited, especially regarding voice loudness control. Previous studies have predominantly focused on pitch modulation, often yielding inconclusive results regarding sensory feedback deficits in the IwPD ((Huang *et al*., 2016). Thus, the neural correlates underlying the impaired regulation of voice loudness in PD so far remains poorly understood.

In the present study, we recruited IwPD and HC participants to investigate whether PD alters ERP responses (P50, N100, and P200) in a task that involved sensory feedback processing of one’s own voice. We used a well-established voice play-back paradigm inspired by Knolle et al. (2019), and examined ERP morphological changes elicited by self-generated versus externally generated voice. Based on previous findings, we hypothesized that IwPD should show a diminished sensory attenuation effect and altered ERP amplitudes, reflecting deficits in predictive sensory processing that are critical for accurate voice modulation.

## 2. Materials and Methods

### 2.1 Participants

We recruited a total of 58 participants, including 30 IwPD and 28 HC. IwPD were approached at the Movement Disorders Unit of Neurology at the Parc Sanitari Sant Joan de DéuMHN-PSSJD in Barcelona. The HCs were recruited from older adult courses at the University of Barcelona and from the family members of IwPD. The experimental protocol included two sessions. In the first session, demographic data (age, sex) were collected from both IwPD and HC, and clinical histories (disease duration, UPDRS-III motor score) were obtained for IwPD. Cognitive and voice assessments were performed by a Speech-Language Pathologist (SLP) specializing in the PD voice to rule out vocal disorders unrelated to PD. The second session involved the EEG experiment. All participants provided written informed consent in accordance with the Declaration of Helsinki and regulations of the Human Research Ethics Committee. The study protocol was approved by the Ethics Committee of the Hospital Parc Sanitari Sant Joan de Déu (CEIm: PIC-27-23).

During the selection process, a specific exclusion criterion for voice quality was used. First, a SLP assessed the overall voice quality of all participants (both HC and those with PD) using a visual analog scale to identify and rule out dysphonia, which is a gold standard of assessment in the voice clinic (Contreras-Ruston et al., 2024). If the voice was found to be dysphonic or there was a diagnosed voice disorder unrelated to PD (e.g., nodules, polyps, functional dysphonia, or other laryngeal lesions), the individual was excluded from the study. This approach ensured that the vocal characteristics analyzed were solely attributable to PD rather than to other voice pathologies.

Second, hearing was assessed in the IwPD and HC groups. This was further verified using a pure-tone detection task involving six frequencies (500, 750, 1000, 1500, 2000, and 4000 Hz) presented at five intensity levels (50, 40, 30, 20, and 10% of the maximum intensity). The task was conducted in a soundproof environment using PsychoPy software v2023.3 (Peirce *et al*., 2019), following a procedure similar to that described by Abur and colleagues (Abur *et al*., 2018; Abur & Stepp, 2020).

Finally, 7 PD participants were excluded due to excessive EEG artifacts, a low score on the Montreal Cognitive Assessment (MoCA) for cognitive impairment (≤26; Nasreddine et al.,2005), or dropout in the second session. In addition, none of the participants had undergone deep brain stimulation or had any other neurological disorders. The final participant sample included 23 IwPD (14 men and 9 women) aged 52 to 83 years (SD = 8.33). All IwPD were diagnosed by a neurologist and met the criteria for early and mid-stage PD (Hoehn & Yahr stages I–III). They were also evaluated using the Unified PD Rating Scale (UPDRS Part III) for speech (see Table 1) in the “ON” state (e.g., antiparkinsonian medications).

**Table 1.** Demographic and Clinical Characteristics of the IwPD.

| Individuals | Sex | Age | Disease Duration (years) | MoCA | H & Y | UPDRS III |
| --- | --- | --- | --- | --- | --- | --- |
| 1 | F | 74 | 17 | 26 | II | 1 |
| 2 | M | 68 | 6 | 26 | I | 2 |
| 3 | M | 64 | 3 | 27 | II | 1 |
| 4 | F | 57 | 5 | 28 | I | 0 |
| 5 | F | 55 | 5 | 27 | I | 1 |
| 6 | M | 52 | 4 | 30 | I | 2 |
| 7 | M | 62 | 20 | 26 | I | 1 |
| 8 | F | 70 | 5 | 28 | II | 1 |
| 9 | M | 52 | 1.2 | 27 | II | 1 |
| 10 | M | 66 | 1 | 27 | II | 1 |
| 11 | M | 56 | 2 | 27 | III | 1 |
| 12 | M | 74 | 2 | 26 | I | 1 |
| 13 | M | 76 | 1.6 | 28 | I | 1 |
| 14 | M | 66 | 5 | 29 | III | 2 |
| 15 | F | 66 | 1 | 27 | I | 1 |
| 16 | F | 58 | 2 | 28 | I | 0 |
| 17 | M | 73 | 1.6 | 27 | II | 2 |
| 18 | M | 60 | 1.6 | 28 | II | 1 |
| 19 | M | 83 | 5 | 26 | III | 3 |
| 20 | F | 67 | 7 | 29 | II | 0 |
| 21 | F | 57 | 8 | 29 | II | 1 |
| 22 | F | 74 | 8 | 29 | II | 1 |
| 23 | M | 64 | 11 | 26 | III | 1 |

Note: M = male; F = female; Disease Duration = Years of diagnosis; H & Y = Hoehn and Yahr stage; MoCA = Montreal Cognitive Assessment score (range: 0–30; $\geq 26$ = normal cognitive level); UPDRS III = Movement Disorder Society-Unified Parkinson's Disease Rating Scale, Part III.

For the HC group, 28 participants were recruited from the general population, of whom five were excluded due to excessive artifacts in EEG signals or dropout, leaving 23 HC (6 men and 17 women) aged between 51 and 70 years (SD = 5.92). In total, 46 participants were included in the study (23 IwPD and 23 HC).

### 2.3 Experimental Design and Procedure

#### 2.3.1 Procedure and Materials

In a first session, we recorded the voices of all the participants in a controlled environment with ambient noise levels below 40 dB. The participants’ tasks included producing three sustained /a/ vowels, each lasting more than three seconds at their habitual loudness and pitch levels, ascending and descending glissandos, and reading aloud ((Patel *et al*., 2018; Rusz *et al*., 2021). The recording equipment included a laptop computer running the Audacity software (© 2023 Audacity Team; https://www.audacityteam.org) and a Karsect HT-9 microphone with a unidirectional polar pattern connected to an external Focusrite audio interface. The microphone was positioned 5 cm away from the participant’s mouth (Patel *et al*., 2018).

In the second session, the most stable of the three vowels was selected as the target stimulus for the EEG experiment. We utilized the playback paradigm described in Knolle et al. (2019), which included three blocked experimental conditions (see Fig 1): auditory-motor condition (AMC), auditory-only condition (AOC), and motor-only condition (MOC). The AOC and AMC conditions involved the presentation of the participant’s own voice at normal loudness and a loudness increase (+15dB). In the AMC, each button press triggered the immediate playback of the participant’s recorded vowel for 0.5 seconds, presented through headphones either at its normal loudness or manipulated by a +15 dB increase. The two types of target stimuli were randomized in the AMC block. In the AOC, participants did not press a button, but were instructed to observe a cross on the screen while attentively listening to recordings of their own voice, as presented in the AMC. In the MOC, participants pressed a button following a visual cue without receiving any auditory feedback.

**FIGURE 1.**
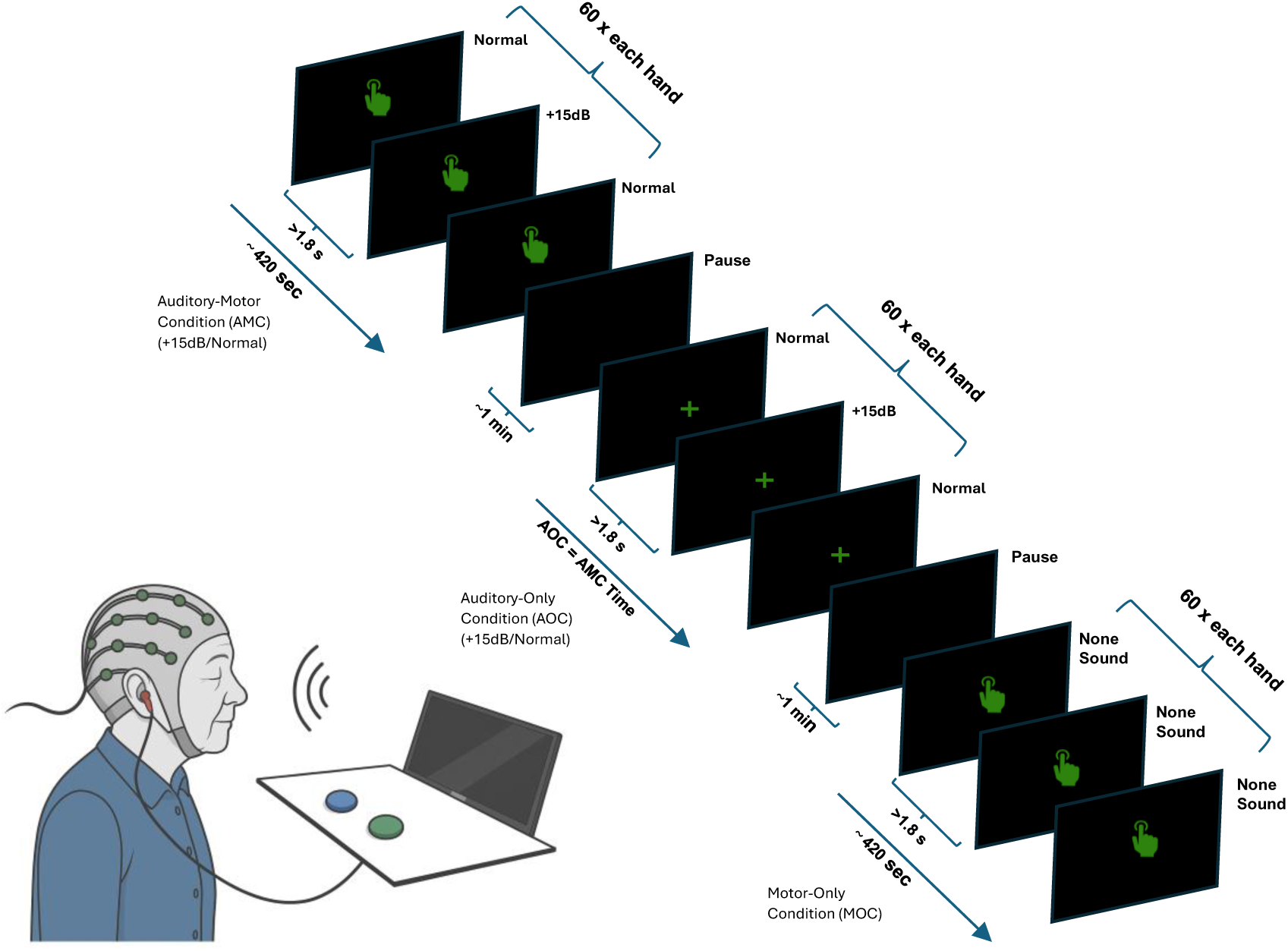
Playback paradigm: In the AMC condition, participants pressed a button at intervals >1.8 s to trigger 420 ms playback of their own voice (normal / +15 dB) in blocks of ∼420 s followed by 1 min rest; the AOC condition uses identical auditory stimuli and timing but without button presses, participants fixated on a central cross, followed by 1 min rest; in the MOC condition participants only press the button at the same cadence (>1.8 s) in ∼420 s blocks with no auditory stimuli, followed by 1 min rest. Each condition was repeated 60 times for each hand.

In both AMC and MOC conditions, participants pressed the button approximately every 2.4 seconds when a fixed visual cue appeared. Button presses within 1.8 seconds (“too early”) were considered as errors. During the experiment, participants completed a total of 60 button presses with their right hand and 60 with their left hand, distributed in two sub-blocks (i.e., AMC-right and AMC-left), conducted in counterbalanced order across participants. The order in which the stimuli were presented in the AOC was the same as that in the AMC for each participant.

Before starting the experiment, participants underwent preliminary training of the paradigm with the same instructions as in the main experiment to fully understand the task and to ensure that the press between each button occurred at an interval greater than 1.8 seconds.

### 2.4 EEG Recording

EEG data were recorded using 64 Ag/AgCl actiCAP-Slim electrodes (BrainAmp, Brain Products) arranged according to the extended 10–20 international system. EEG signals were acquired using two BrainAmp amplifiers, each sampling at 250 Hz. The electrode impedance was maintained below 10 kΩ throughout the recording. The FCz electrode served as an online reference, whereas the Fpz electrode was used as the ground electrode. Data acquisition and storage were performed using Brain Vision Analyzer 2 (version 2.1.1.2516).

### 2.5 Data Analysis

#### 2.5.1 EEG Preprocessing

The EEG data were preprocessed in MATLAB using the EEGLAB toolbox (Delorme & Makeig, 2004). First, a notch filter (*lineFilter*) was applied to remove line noise, followed by a band-pass filter (0.05–35 Hz) using a finite impulse response (FIR) design with a Kaiser window, ensuring a linear-phase response to minimize temporal distortions. Channels with poor signal quality were then automatically identified and removed using (*pop_clean_rawdata*), based on the FlatlineCriterion (≥5 s without variation), ChannelCriterion (correlation <0.8), and Distance = ’Euclidean’ (detection of atypical channels). Removed channels were subsequently interpolated and the data were re-referenced to the mastoid electrodes. To ensure precise synchronization between button presses and stimulus presentation, event latencies were corrected by adjusting markers according to the measured delay. A Robotic Key Actuator (RKA) from The Black Box

ToolKit (Version 2, Rev. RC2) (Plant, 2013) was used to determine and compensate for the mechanical latency of the actuator, allowing for a precise alignment between participant responses and stimulus onset. Relevant triggers for each experimental condition were then identified, and data were segmented into epochs from –2 s to +0.5 s relative to these events.

The epoched data were subjected to ICA using (*pop_runica*) in extended mode to decompose the signals into independent components. These components were automatically classified using ICLabel, marking as artifacts those with a probability ≥80% for ocular activity and ≥60% for muscular activity (*pop_icflag*). Artifacts were removed with (*pop_subcomp*), and the EEG signal was reconstructed from the remaining components to retain genuine brain activity. Finally, an additional cleaning step was applied to remove epochs with amplitudes exceeding ±100 µV using (*pop_eegthresh*) for automatic detection. Additionally, (*pop_jointprob*) was used to exclude epochs with extreme values in joint probability and power distribution, with adaptive thresholds set based on intersubject variability. A final visual inspection ensured the effective removal of residual artifacts and overall quality of the preprocessed data.

#### 2.5.1 Event- Related Analyses

After preprocessing and artifact rejection, the epochs were averaged separately for each experimental condition to obtain the ERP waveforms. ERP data were processed by averaging electrodes from regions of interest (ROIs), including the frontal (F1, Fz, F2), frontocentral (FC1, FCz, FC2), and central (C1, Cz, C2, C3, C4) areas, following previously established criteria (Rosburg et al., 2008; Knolle et al., 2019; Pinheiro et al., 2019). Three experimental conditions were included: AMC with two different sound loudness levels (normal and +15 dB), AOC with the same two loudness levels, and MOC.

Baseline correction was applied using an interval of -100 ms to 0 ms before the stimulus. To isolate the neural responses to sounds from those generated by pressing the button, we subtracted the MOC from the AMC trials. This correction removed the contribution of motor-related activity associated with movement execution, ensuring that the resulting evoked potentials primarily reflected the auditory response to self-generated sounds in the AMC, and thus could be compared with those from the AOC. This procedure follows previous sensory suppression playback paradigms (e.g., Knolle et al., 2013), which allow the evaluation of the impact of sensory feedback processing. For statistical analysis, time windows were defined based on previous studies (Huang *et al*., 2016; Knolle *et al*., 2019; Li *et al*., 2021) and a grand average across all combined conditions (Murray et al., 2008; Luck, 2014). The peaks of the P50 (40–75 ms), N100 (90–150 ms), and P200 (190–250 ms) components were extracted, along with their mean amplitudes, variances, peak amplitudes, and latencies. Finally, the number of trials per participant was equalized for each ERP component using random sampling (*randperm*) to ensure a comparable signal-to-noise ratio across groups and conditions and to maintain an equivalent amount of data for between-subject comparisons.

#### 2.5.2 Statistical Analyses

All analyses were conducted using R (R Core Team, 2024) within the RStudio environment (version 2024.12.0-467). To examine individual differences in response variability, multilevel models (multilevel analysis) were employed. This approach is particularly useful in studies investigating response stability at both intra- and inter-group levels (Volpert-Esmond *et al*., 2021):

1. Relaxes the assumption of homoscedasticity and allows the examination of differences in response variability.
2. It includes random effects that capture inter-individual heterogeneity.
3. It improves statistical efficiency and reduces the likelihood of Type I and Type II errors. In the present study, a restricted equality model was used to analyze the amplitude and latency of all ERP components by applying identical parameters to each component in the independent models. This approach optimized parsimonious analysis, prevented overfitting, and ensured a more robust interpretation of the results.

To evaluate the effects of group (HC vs. PD), condition (AMC vs. AOC), and Stimulus loudness (Normal vs. +15 dB) on P50, N100, and P200 amplitude and latency, multilevel location-scale models were combined with linear mixed-effects models (LMMs).

The models used were as follows:

lmer(varOI ∼ Group + Condition + Loudness + Group:Loudness + Condition:Loudness + Group:Condition:Loudness + (1 | Participants)), Where varOI (variable of interest) represents either the amplitude or latency of the ERP components. To this end, we removed the Group × Condition interaction, as both groups were expected to show a condition effect based on our hypothesis. We then applied stepwise backward elimination based on coefficient p-values (α > 0.05), removing non-significant high-order interactions (Kuznetsova *et al*., 2017).

Fixed-effect estimates, 95% Wald confidence intervals, and p-values were calculated. Random-effect variances were used to calculate the intraclass correlation coefficient (ICC). Model performance was assessed using marginal (R²_marginal) and conditional (R²_conditional) R² values, representing the variance explained by the fixed effects and the full model, respectively.

To decompose the significant interaction, follow-up simple-effect analyses were performed separately within each group (HC, PD) for both amplitude and latency. Post-hoc pairwise comparisons of the estimated marginal means were conducted with Tukey’s adjustment for multiple comparisons. All models were estimated using the *lme4* package (Bates *et al*., 2015) and *lmerTest* (Kuznetsova *et al*., 2017) to obtain *p*-values based on Satterthwaite approximation. The *emmeans* package (Lenth, 2021) was used for post-hoc comparisons, and effect size (Ben-Shachar *et al*., 2020) for effect size estimation. Tabular summaries of the fixed effects were generated using *sjPlot*. Data visualization was carried out using *ggplot2* (Wickham, 2016). Statistical significance was defined as p < .05.

## 3. Results

### 3.1. Multilevel Analysis

#### 3.1.1. P50

For the P50 amplitude, we started with a full model including all main effects, two-way interactions, (Group × Loudness) and (Condition × Loudness), and three-way interaction (Group × Condition × Loudness). In a stepwise fashion, we removed non-significant interactions and re- fitted the model. As none of the interaction terms were significant, the final model included only the main effects.

The final model revealed a significant main effect of Condition, *F*(1, 6208) = 24.22, *p* < .001, indicating higher P50 amplitudes in the AOC condition than in the AMC condition (Estimate for AMC vs. AOC = –0.88, 95% CI [–1.23, –0.53], *p* < .001). Furthermore, the model showed a significant main effect of Loudness: *F*(1, 6208) = 5.05, *p* = .025 (Estimate = –0.40, 95% CI [– 0.75, –0.05]), suggesting that stimulus loudness modulated the P50 amplitude. By contrast, no effect of Group (IwPD vs. Controls) was found, *F*(1, 44) = 0.01, *p* = .904 (Estimate = –0.04, 95% CI [–0.69, 0.61]). In the final model, inter-individual variability at the participant level was moderate (τ₀₀ = 0.91) and the residual variance was 50.16. The marginal R² (variance explained by the fixed effects alone) was 0.005, whereas the conditional R² (variance explained by both fixed and random effects) was 0.022.

For P50 latency, the final model included only the main effects of Group, Condition, and Loudness, as neither the three-way nor two-way interactions were significant. A significant main effect of condition emerged, F(1, 6208) = 53.70, p < .001, indicating shorter latencies in the AMC condition compared with AOC (Estimate for AMC vs. AOC = –0.00, 95% CI [–0.00, –0.00], p < .001). The main effects of Group F(1, 44) = 3.09, p = .086 (Estimate = –0.00, 95% CI [–0.00, 0.00]) and Loudness were not significant, F(1, 6208) = 1.17, p = .280 (Estimate = 0.00, 95% CI [–0.00, 0.00]). The model indicated very low interindividual variability (τ₀₀ = 6.886 × 10⁻⁷) and a residual variance of 1.381 × 10⁻⁴. The marginal R² for latency was 0.009 and the conditional R² was 0.014, indicating that only a small portion of the total variance in latency was explained by the combined fixed and random effects.

#### 3.1.2. N100

For the N100 amplitude, the initial model included all main effects, two-way interactions, and three-way interactions between Group, Condition, and Loudness. The three-way interaction between these factors reached significance and was therefore maintained.

For N100 amplitude, the effect of Condition was significant (β = 4.06, 95% CI [3.57, 4.55], p < .001), as was the main effect of Loudness (β = 0.78, 95% CI [0.14, 1.43], p = .018). A significant Group × Condition × Loudness interaction was observed (β = –0.99, 95% CI [–1.97, –0.01], p = .048). The random-intercept variance was τ₀₀ = 4.71 (ICC = 0.09), with marginal R² = 0.054 and conditional R² = 0.137.

Post-hoc comparisons revealed that louder stimuli significantly increased in N100 amplitude (i.e., elicited greater negativity) in the AOC condition for HC (estimate = –0.83, SE = 0.353, p = .0185), with no effect in AMC. In IwPD, an even larger increase in amplitude (greater negativity) was observed in AOC (estimate = –1.34, SE = 0.355, p = .0002), with no significant effect of loudness in the AMC condition (p > .05).

For N100 latency, significant main effects of Condition (β = –0.01, 95% CI [–0.01, –0.01], p < .001) and Loudness (β = 0.00, 95% CI [0.00, 0.00], p = .030) were found, indicating shorter latencies in the AMC condition and in response to louder stimuli. No significant interactions were observed in the final model. Inter-individual variability was low (τ₀₀ = 1.47 × 10⁻⁵), with an ICC of 0.05. The marginal R² was 0.034 and the conditional R² was 0.080 (see Table 2).

**Table 2.**
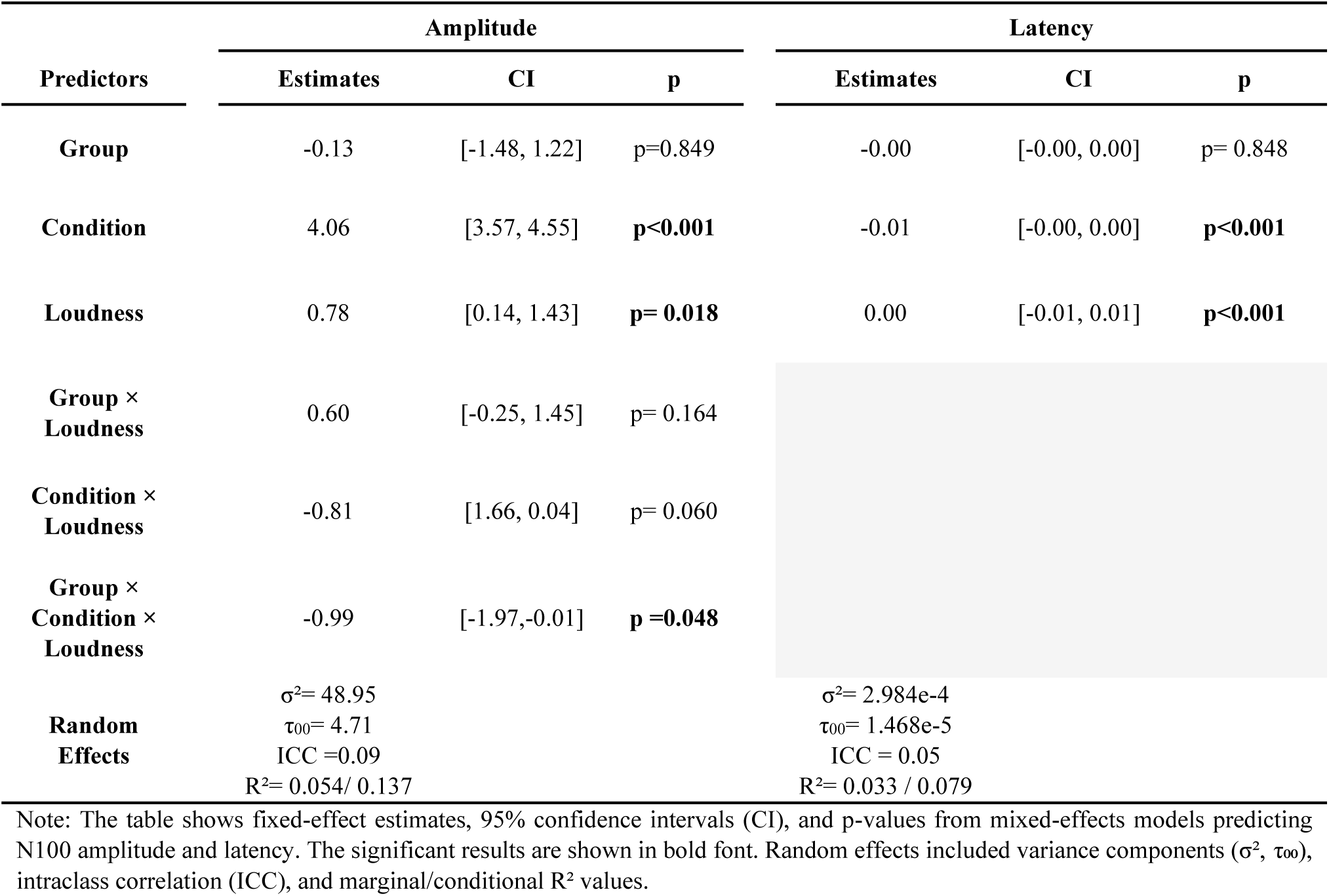
Effects of Group, Condition, and Loudness on N100 Amplitude and Latency.

Post-hoc comparisons revealed that both groups showed reduced latencies in response to louder stimuli across conditions. In the HC group, latency was significantly reduced in the AMC (p = .0006) and AOC (p = .0063). In IwPD, significant latency reductions were also found in AMC (p = .0050) and only marginally in AOC (p = .0507) (see Fig. 2 and Table 3).

**FIGURE 2.**
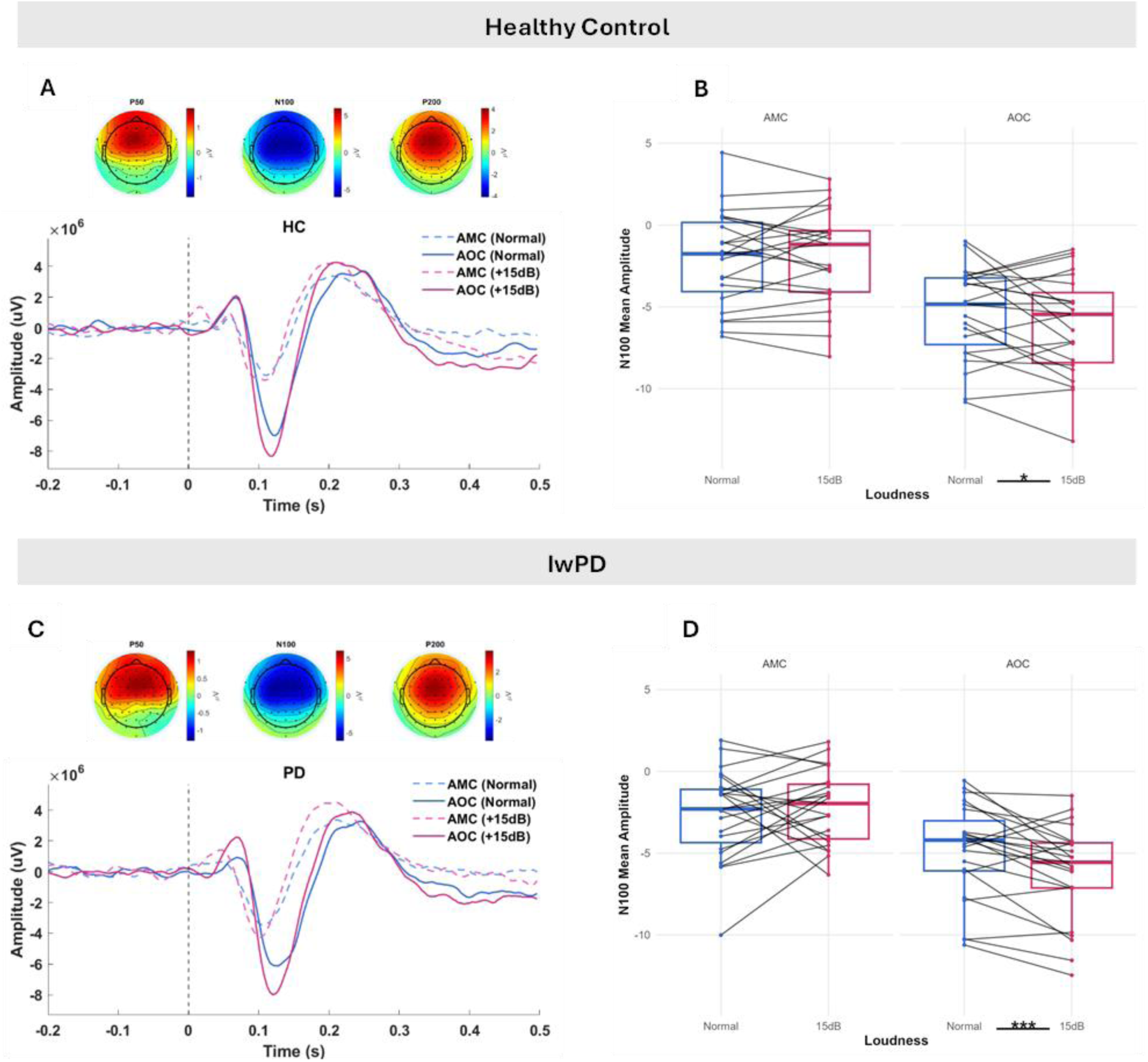
Event-related potentials (ERPs) across conditions and groups. A–C. Grand average ERP waveforms and scalp topographies of HC (panel A) and IwPD (panel C). ERPs are time-locked to stimulus onset (vertical dashed line at 0 s) and displayed from –0.2 to 0.5 seconds. The y-axis shows the amplitude in microvolts (µV). Four conditions were plotted: AMC with normal loudness (solid blue), AMC with increased loudness (+15 dB; solid magenta), AOC with normal loudness (dashed blue), and AOC with increased loudness (dashed magenta). The scalp topographies above each ERP reflect the spatial distribution of the voltage at the peak latency of the P50, N100, and P200 components. Waveforms were averaged over the frontocentral region of interest. B–D. N100 mean amplitude (µV) for each participant across loudness levels (Normal vs. +15 dB) and auditory conditions (AMC vs. AOC), for the HC (panel B) and IwPD (panel D) groups. Each black line represents a single participant. Boxplots display group-level distributions, and dots are colored by loudness (blue = Normal, magenta = +15 dB). A significant loudness effect was observed in the AOC condition for both groups (*p <.05, HC; ***p <.001, PD).

**Table 3.** Post-hoc comparisons of Condition, and Loudness on N100 Amplitude.

| **Loudness Effect:**<br>**Normal – 15dB** |  | **Amplitude** |  |
| --- | --- | --- | --- |
| **AMC** |  | **Estimate** | **p-value** |
| N100 HC |  | 0.075 | 0.830 |
| N100 PD |  | 0.368 | 0.299 |
| **AOC** |  |  |  |
| N100 HC |  | -0.831 | **p=0.018** |
| N100 PD |  | -1.342 | **p<0.001** |
| **Condition Effect:**<br>**AMC – AOC** |  |  |  |
| **Normal Loudness** |  | **Estimate** | **p-value** |
| N100 HC |  | 3.24 | **p <0.0001** |
| N100 PD |  | 2.25 | **p<0.0001** |
| **+15 dB** |  |  |  |
| N100 HC |  | 4.15 | **p <0.0001** |
| N100 PD |  | 3.96 | **p<0.0001** |

Note: The table presents post-hoc pairwise comparisons of Loudness (Normal vs +15 dB) and Condition (AMC vs AOC) for the N100 responses in healthy controls (HC) and individuals with Parkinson's disease (PD). Values are estimated differences (Estimate) with associated p-values; significant comparisons ( $p < .05$ ) are indicated in bold.

#### 3.1.3. P200

Regarding the P200 component, no significant interactions were found. The final model included only the main effects of the Group, Condition, and Loudness. A significant main effect of Loudness was observed for amplitude (β = –0.62, 95% CI [–0.98, –0.26], p = .001), indicating that louder stimuli elicited an increased P200 amplitude. No significant effects were found for Group or Condition. The model showed moderate inter-individual variability (τ₀₀ = 3.91), with an intraclass correlation coefficient (ICC) of 0.07. The marginal R² was 0.002, and the conditional R² was 0.069.

For P200 latency, significant main effects were found for both Condition (β = –0.00, 95% CI [– 0.00, –0.00], p < .001) and Loudness (β = 0.00, 95% CI [0.00, 0.00], p < .001), reflecting shorter latencies in the AMC condition and louder stimuli. No significant effect of Group was observed. Inter-individual variability was minimal (τ₀₀ = 8.21 × 10⁻⁶), with ICC = 0.02. The marginal R² was 0.004 and the conditional R² was 0.026.

## 4. Discussion

Auditory sensory feedback plays a critical role in voice regulation, allowing the central nervous system to monitor and adjust parameters such as loudness, pitch, and articulation in real time (Behroozmand *et al*., 2016). In IwPD, this regulatory mechanism is likely impaired, contributing to hypophonia and the vocal monotony characteristics of hypokinetic dysarthria (Liu *et al*., 2012; Huang *et al*., 2016). One way to investigate the integrity of this feedback system is through EEG, specifically by examining the ERPs elicited during sensory feedback in voice tasks. The morphology and amplitude of early auditory ERPs—namely the P50, N100, and P200 components—are thought to reflect the brain’s ability to detect and process self-generated voice sounds. Therefore, altered ERP responses may serve as physiological markers of disrupted sensory feedback in IwPD.

Modulations in P50 amplitude were previously associated with sensory gating, attenuating responses to repeated stimuli, and protecting the auditory system from overstimulation (Brinkmann *et al*., 2021). In this study, we found that the P50 response was significantly larger in the AOC condition (self-voice presented externally) than in the AMC condition (self-generated voice) in both groups, suggesting increased activation in response to external auditory stimuli, even when the stimulus was acoustically recognizable, such as one’s own voice. Additionally, there was a minor but significant loudness effect on the P50 amplitude, which increased with a (+15 dB) loudness increment. However, no significant differences emerged when comparing the groups, nor did we find any interaction between stimulus loudness and group. This observation may indicate that, at least in the current experimental paradigm, no significant differences in the P50 amplitude were found between PD and HC, indicating comparable early sensory responses. These discrepancies may reflect the engagement of predictive sensory-feedback processes when individuals hear their one own voice, processes not recruited by meaningless external sounds. In our paradigm, such voice specific sensory-feedback processing could contribute to preserved habituation, in contrast to the reduced habituation reported for non-vocal stimuli in PD ((De Groote *et al*., 2020; Hassin-Baer *et al*., 2022).

Regarding the P50 latency, both the PD and control groups showed significant differences between AOC and AMC conditions, consistent with the idea that motor preparation facilitates the processing of self-generated stimuli (Pinheiro *et al*., 2019). Although there was a trend toward a Group × Condition interaction, this did not reach statistical significance. The P50 amplitude itself did not differ between groups; however, its modulation by stimulus origin and loudness suggests that P50 sensory gating remains intact in IwPD, with PD-related impairments potentially emerging only in later ERP components such as N100 and P200 (Hirano *et al*., 2008).

The N100 component modulation in response to self-generated sounds reflects auditory cortical suppression, indicating attenuation of expected sensory feedback (Knolle *et al*., 2013b, 2019). Previous studies have reported reduced or absent N100 suppression effects in IwPD (Huang *et al*., 2016; Mollaei *et al*., 2016), suggesting impairments in sensory feedback processing (Contreras-Ruston *et al*., 2025). Here, a significant interaction between the group, condition, and loudness in the N100 response was observed. This interaction and follow-up post-hoc analyses indicated that IwPD and controls responded differently to sound loudness changes depending on whether the voice was self-generated or externally produced. This observation aligns well with findings from Brajot et al. (2016), who noted a sensory feedback impairments in IwPD specifically during loudness regulation of the self-generated voice, showing different response patterns in iwPD and HC in altered auditory feedback conditions (Brajot *et al*., 2016). Specifically, in the current study, increasing the loudness of externally generated stimuli (i.e., participants’ own pre-recorded voices), the N100 amplitude was significantly more negative only in the AOC condition in both groups. This effect was more pronounced in IwPD, suggesting hyper-reactivity in response to external sensations with increased loudness.

Electrophysiological evidence has shown that IwPD show larger-amplitude FFRs to speech stimuli, reflecting enhanced neural encoding of pitch and sound loudness during early processing stages (Mollaei *et al*., 2022). Similarly, unmedicated IwPD display abnormally large auditory brain oscillations in the delta, theta, and alpha bands, particularly in the parietal regions, suggesting a breakdown of inhibitory control over cortical sensory feedback processing (Güdücü *et al*., 2019). Taken together, these findings are consistent with altered sensory feedback processing in basal ganglia cortical circuits as a possible contributing factor to IwPD’s increased sensitivity to external sounds, although other process, e.g., attentional modulation (Praamstra & Oostenveld, 2003) or inhibition (Ziri *et al*., 2024), cannot be ruled out.

However, from the perspective of forward models, evidence remains unclear regarding loudness manipulation on the N100, as a study using pure tones of varying intensities in IwPD did not find significant differences in either the amplitude or latency of the N100 compared to controls (De Keyser *et al*., 2021). Whether the perception of a self-generated voice is equivalent to that of pure tone remains an open question. Pinheiro et al., (2018) directly compared, in individuals with schizophrenia and controls, self-initiated tones and voices in nonclinical voice-hearers with low and high hallucination-proneness (HP) using a button-press paradigm. Low-HP participants replicated the classic N100 suppression effect for both stimuli, but high-HP participants showed a reversal of this suppression (an enhanced N100 amplitude) only for self-generated voices, while their responses to tones remained unchanged. In contrast, patients with schizophrenia exhibit reduced sensory attenuation of the N100 to self-initiated auditory stimuli, affecting both tones and voices and reflecting generalized forward model dysfunction. Since De Keyser et al. (2021) did not evaluate self-generated voices, it remains to be investigated whether N100 suppression to one’s own voice in PD is preserved or follows a different pattern.

Here, both groups showed an N100 suppression effect, but in IwPD the response to external stimuli was more enhanced than in HC when loudness changed. This suggests that although cortical mechanisms of suppression remain partially preserved in PD, processing of externally generated voice stimuli may be impaired. Specifically, IwPD responded differently to externally generated sounds than HCs, an observation consistent with previous findings (Huang *et al*., 2016; Mollaei *et al*., 2016). However, given the lack of a significant difference in the AMC and the absence of a control condition using pure tones, we cannot conclusively attribute this difference to impaired sensory feedback processing per se. Instead, these results point to a possible alteration in the way IwPDs perceive the loudness of their own externally presented voice sounds.

This finding partly supports our hypothesis that IwPDs have difficulty processing auditory feedback, consistent with preserved N100 attenuation during self-generated voice (Huang *et al*., 2016) and impaired motor prediction mechanisms (Mollaei *et al*., 2016). The enhanced N100 amplitude observed in IwPD during AOC may reflect an impaired ability to modulate auditory cortical responses to externally presented self-voice stimuli. The reduced attenuation observed here could reflect dysfunction in the predictive mechanisms responsible for auditory stimulus processing, partially aligning with previous reports of impaired sensory suppression during self-generated voice in PD (Railo *et al*., 2020). However, Railo et al. specifically reported reduced suppression during active (i.e., auditory-motor) and passive self-voicing (i.e., auditory-only) conditions. Therefore, caution is required in interpreting our results as evidence of impaired sensory suppression, given that our paradigm shows similar AMC suppression in both groups, but a difference in the AOC responses. Importantly, in Railo and colleagues (2020), there was no manipulation of the loudness of one’s own voice in both the active and passive conditions. Therefore, our study delivers an important novel aspect, the perception of one’s own voice loudness, considering that a hallmark symptom of IwPD is a voice with reduced loudness that they often fail to self-perceive. Regarding N100 latency, there was no significant three-way interaction, but significant main effects of condition and loudness: latency differed between AMC (self-generated) and AOC (externally) generated voice conditions, and between the normal and +15 dB loudness levels.

P200 amplitude modulations have been associated with auditory–motor integration and compensatory responses to altered auditory feedback (Huang *et al*., 2019). In PD, an increased P200 amplitude in response to unexpected auditory signals during speech production were observed and interpreted as a heightened corrective effort or cortical compensation when feedforward control is insufficient to maintain voice stability (Liu *et al*., 2012; Chen *et al*., 2013; Huang *et al*., 2016). This feedforward control is thought to be mediated mainly by non-cortical structures such as the basal ganglia and cerebellum (Hickok *et al*., 2011; Tourville & Guenther, 2011; Parrell & Houde, 2019). Here, no significant group differences were found in this ERP component, but there was a significant effect of sound loudness; louder stimuli reduced the P200 amplitude and slightly modulated its latency. This may be explained by a stronger and quicker electrophysiological response to louder sounds, which demand fewer cortical resources (Covington & Polich, 1996).

The present findings provide some evidence in favor of the hypothesis that it is not sensory feedback processing per se, but rather the processing of one’s own voice presented externally during passive listening, that may contribute to voice deficits in IwPD. Unlike previous suppression studies, where reduced suppression typically arises from altered auditory motor processing, our findings indicate that the N100 suppression in PD was driven by changes in the AOC condition. This suggests a distinct alteration in the perceptual processing of externally presented voice stimuli, rather than a deficit specific to self-generated auditory-motor feedback. The limited modulation of the P50 and P200 components between groups indicates that the most significant differences emerge at the early sensory feedback processing stage, as indexed by N100 modulation Nevertheless, previous studies have reported heterogeneous results regarding ERP components in PD, including an increased P200 amplitude, interpreted as a compensatory mechanism when the feedforward control system is compromised (Huang et al., 2016; Knolle et al., 2019). Such variability across studies might be attributable to differences in experimental paradigms, measures used (e.g., voice loudness or pitch), and clinical heterogeneity inherent to PD (Contreras-Ruston et al., 2025).

Altogether, the present study provides novel evidence on voice production and sensory feedback processing in PD. However, some limitations should be noted. First, our relatively small and narrowly defined cohort, 23 IwPD at early-to-moderate stages (I–III) under a single medication regimen, plus 23 matched healthy controls, limits the extent to which these findings can be extrapolated to the broader PD population. In particular, patients at more advanced stages, on different treatment protocols, of varying ages or cognitive status, may exhibit distinct patterns of N100 modulation. Future work with larger, more diverse samples spanning later disease stages, multiple medication conditions, and broader demographic ranges is needed to confirm the generalizability of our results. In addition, it was not possible to optimally match participants by age and sex, and the duration of Parkinson’s diagnosis varied among the individuals. All PD participants were assessed in the “ON” state of their medication, preventing assessment of how different dopaminergic levels might affect ERPs. Finally, although our button-press paradigm using isolated sustained vowels lacks the communicative richness of spontaneous speech or reading aloud, its highly controlled design is essential for isolating specific neural processes.

Indeed, previous work in healthy subjects using similar playback designs has demonstrated robust N100 suppression for self-initiated versus externally generated sounds (Pinheiro *et al*., 2019, 2020). In contrast, we observed reduced N100 suppression only in the AOC condition in IwPD. This divergence underscores the need for future studies to manipulate the delay between motor action and playback in comparable self-initiated paradigms (Korzyukov *et al*., 2017) and incorporate more naturalistic stimuli, such as words or phrases, to better approximate spontaneous speech in PD and extend these findings to everyday communication.

Despite these limitations, these results encourage future research on the role of sensory feedback processing in PD speech production. We consider that this approach could be further complemented by neuroimaging methods, including EEG-based functional-connectivity analyses (e.g., coherence, phase-locking), to better examine functional connectivity patterns during speech tasks. Basal ganglia structures, particularly the putamen and external globus pallidus, have been shown to increase their activation in response to stimuli with higher loudness, while the anterior caudate nucleus exhibits an inverse relationship with perceived loudness (Grahovac & Klostermann, 2023). This can be linked to exaggerated responses to the loudness of one’s own voice. For example, Rektorova et al. (2012) documented altered connectivity between the cortical and subcortical regions, supporting the hypothesis of altered sensory integration processes. Furthermore, the implementation of paradigms that combine speech-induced suppression with motor-related potentials, such as priming potentials (Tremblay & Sato, 2024), could provide a complementary characterization of how motor priming and auditory feedback interact during speech production in the IwPD. Responses occurring after 300 ms, reflecting the evaluation and updating of internal models following unexpected auditory feedback, reveal how IwPD detect and integrate mismatches in their self-generated voice. This idea is supported by Suchý et al., (2023), who demonstrated in healthy controls that conscious detection of pitch-shifted feedback errors elicits a robust late positive ERP (500–700 ms), correlating with compensatory vocal responses and involving temporal, frontal, and parietal networks.

From a clinical perspective, the current findings highlight the need to develop tools that enable visualization and quantification of voice self-perception in IwPD, such as real-time auditory feedback interfaces. Furthermore, both auditory assessments (Castillo-Allendes *et al*., 2021) and voice therapies or speech technologies targeting this population should explicitly incorporate training strategies that address auditory feedback. Focusing on these sensory processes could enhance the regulation of vocal loudness and, in turn, improve everyday communication in IwPD.

## 7. Conclusion

IwPD displayed a clear hypersensitivity to loudness changes when they heard their own voice as an external stimulus. This indicates a dysfunction in monitoring auditory sensory input from one’s own voice. Specifically, IwPD showed an exaggerated N100 amplification when generating accurate sensory predictions of loudness for playback of their own vocalizations. Thus, the current findings reveal an early deficit in auditory self-monitoring; whether this dysfunction extends to other auditory inputs remains to be determined; this effect might explain less voice loudness in PD, beyond a simple failure of correcting feedback in voice production. From the clinical point of view, this underscores the importance of incorporating rehabilitation strategies that emphasize auditory sensory process and recalibrate how the IwPD perceives their own voice, increasing the loudness of the voice, and mitigate the communicative impact of hypophonia in these individuals.

## Acknowledgment

The authors wish to thank the participants with Parkinson’s disease from Parc Sanitari Sant Joan de Déu for their participation in this study. We also gratefully acknowledge Gemma Colomé for her assistance in recruiting participants and Manuel Moreno for his help in preparing the experiment.

## Data availability statement

The data used here cannot be used for patient data protection reasons.

